# Noxious stimulation induces self-protective behaviour in bumblebees

**DOI:** 10.1101/2024.01.15.575734

**Authors:** Matilda Gibbons, Elisa Pasquini, Amelia Kowalewska, Eva Read, Sam Gibson, Andrew Crump, Cwyn Solvi, Elisabetta Versace, Lars Chittka

## Abstract

Self-grooming directed towards a noxiously-stimulated body part is one indicator that an animal may feel pain. In insects, the lack of evidence for such behaviour has been widely argued to reflect the absence of pain experiences. Here, we tested whether bumblebees (*Bombus terrestris*) selectively groom one of their antennae that was touched with a noxiously-heated (65 □C) probe. In the first two minutes after being touched with the noxiously-heated probe, bees groomed their touched antenna a) more than their untouched antenna, b) more than bees that were touched on the antenna with an unheated probe, and c) more than control (untouched) bees groomed either of their antennae. Our results clearly show that bumblebees can direct grooming towards a site of noxious stimulation. Our findings thus refute arguments that claim that insects do not feel pain because of their lack of displaying this behaviour.

## Background

Nociception is the detection and processing of noxious stimuli [1], whilst pain is the negative subjective feeling that, at least in humans, typically accompanies nociception [2]. Nociception can be identified from neural activity within nociceptive circuits [3] or from the performance of behaviours that require nociceptive circuits [4]. The subjective experience of pain, however, is harder to identify [5]. Even the gold-standard evidence for the subjective experience of pain in humans, verbal self-report, can be unreliable [6,7]. This issue is amplified in other animal species because they cannot verbally describe their pain. However, sets of neural and behavioural indicators to evaluate pain have been developed that attempt to assess whether an animal has the neural capacity and behavioural ability to feel pain [8– 10].

One behavioural indicator of pain is self-protective behaviour directed towards a site of noxious stimulation [8–11]. Examples include tending to, guarding, grooming, and rubbing a noxiously stimulated body part. Consider, for example, the human response to a stubbed toe or a bump on the head to ‘rub it better’, thereby reducing the feeling of pain [12,13]. This self-touch reduces pain, and not just nociception, since pain can be affected with no effect on nociceptive processing [14,15]. An example of this was demonstrated using the thermal grill illusion, where some fingers are warmed while the other fingers are cooled, and, in doing so, the person perceives noxious heat; self-touch (touching their other hand with the affected hand) reduced perception of heat by 64% compared to controls [14]. This is one reason that the Checklist of Nonverbal Pain Indicators in humans includes ‘massaging or clutching the affected area’ [16]. Therefore, the presence of this behaviour is used to indicate that the individual is feeling pain, because it suggests that the person might both be internally representing the bodily location of the aversive stimulus and trying to reduce the pain [17].

Self-protective behaviour is also observed in other vertebrates. For example, rats (*Rattus norvegicus*) rub their face more after their face is injected with a noxious substance [18]. Further, some bird species have been observed grooming a limb that was injected with a noxious substance (e.g. *Pyrrhura molinae*: 19). There are similar findings in fish (*Oncorhynchus mykiss*), where the fish rubbed an area that was treated with a noxious injection into the gravel and the sides of their tank [20]. In these studies, the grooming of the noxiously-stimulated area was taken as evidence for the presence of pain in these non-human animals.

Some invertebrates have also been observed performing self-protective behaviour, in the form of grooming a noxiously-stimulated site. For example, injecting formalin into the claw of Asian shore crabs (*Hemigrapsus sanguines*) induced rubbing of the affected claw [21]. Similarly, applying acetic acid to various body parts induces grooming or scratching in shore crabs (*Carcinus maenas*) [22], prawns (*Palaemon elegans*) [23], cuttlefish (*Sepia pharoaensis*) [24] and octopuses (*Octopus bocki*) [25]. The latter also respond with grooming to an area on their arm that was crushed with forceps for up to 20 seconds [26].

With regards to insects, however, there are no quantitative studies of self-protective behaviour (such as grooming) towards a noxiously-stimulated site [27]. In addition, anecdotal reports claim that insects do not protect their injury sites, and that insects continue to walk, feed, and mate normally after injury [28; 29]. These reports are often cited as evidence against insects experiencing pain [30–32].

Insects are known to self-groom in non-noxious contexts, e.g. during general cleaning [33], and when removing dust particles (e.g. in the *Blattella germanica* German cockroach [34]), pollen grains (e.g. in bees: 35) and parasites such as mites (e.g. in honeybees, *Apis mellifera*: 27). After noxious stimulation, insects may also generally groom more or change their grooming pattern. For example, after having their antenna amputated, red mason bees (*Osmia bicornis*) wipe, or groom, the head and body, although no site-specific measurements were found/taken, nor was there a non-noxious control to compare to [37]. There are also reports that hint insects may self-groom noxiously-stimulated sites, although such reports have not yet been supported by quantitative or statistical analyses [38]. When pinched on the abdominal proleg, *Manduca sexta* moth larvae reportedly turned their heads to the wound, and repeatedly touched the area with their mouthparts, but this behaviour was not measured or compared to a control [39]. *Periplaneta americana* cockroaches appeared to groom their wounds following an abdominal puncture but, again, this behaviour was not measured or compared to a control [40]. Since these observations were not supported by quantitative measurements or analyses [38], a robust, experimental assessment of grooming behaviour in response to noxious stimuli, and as an indicator of pain in insects, is required.

In this study, we tested whether *Bombus terrestris* bumblebees selectively groom a noxiously-stimulated antenna. For each bumblebee, we either briefly touched one antenna with a noxious stimulus (a 65°C heat probe), or a non-noxious tactile stimulus (an unheated probe), or we did not touch either antenna (control). We recorded grooming behaviour on both antennae for 25 minutes.

If bees direct self-grooming towards a site of noxious stimulation, we would predict more grooming on the noxiously-stimulated antenna than the other antenna. We would not expect this difference in bees touched with an unheated probe, nor by bees that were not touched.

## Methods

### Ethics

The UK does not regulate insect welfare in research. Nonetheless, we followed the 3Rs principles [41] in our experimental design and husbandry. In this vein, although some noxious stimulation is required to study pain, we chose a temperature that, when brief, has no long-term effects on the bees (65°C; based on [35]). We also used a power analysis to estimate the minimum required sample size (estimated sample size = 80; alpha: 0.05; power: 80%). According to current best practice, we have followed the ARRIVE guidelines for reporting this research [43].

### Animals and Housing

We used 82 bees from seven bumblebee colonies (standard hives from Biobest Group, Belgium). The bees were housed in ventilated wooden boxes (56×16×11cm; see Figure 1). Each box comprised four sections, arranged linearly and connected by 1cm-diameter holes. At one end was the section containing the nest, which was covered with plywood. The section at the opposite end contained a 35 ml cylindrical feeder (74.5×31 mm), which dispensed Biogluc sugar solution *ad libitum* (Biobest group, Belgium). To access the food source, the bees had to cross the middle two sections. The middle section adjacent to the feeding section was the observation box during the testing period. The floor of both middle sections was covered with a thin layer of cat litter (Catsan Hygiene Plus, Mars Inc, USA) to absorb waste and debris. Each colony received 7g of pollen (Natupol Pollen, Koppert Biological Systems) every two days, and the laboratory was maintained at 23° C.

**Figure 1.**
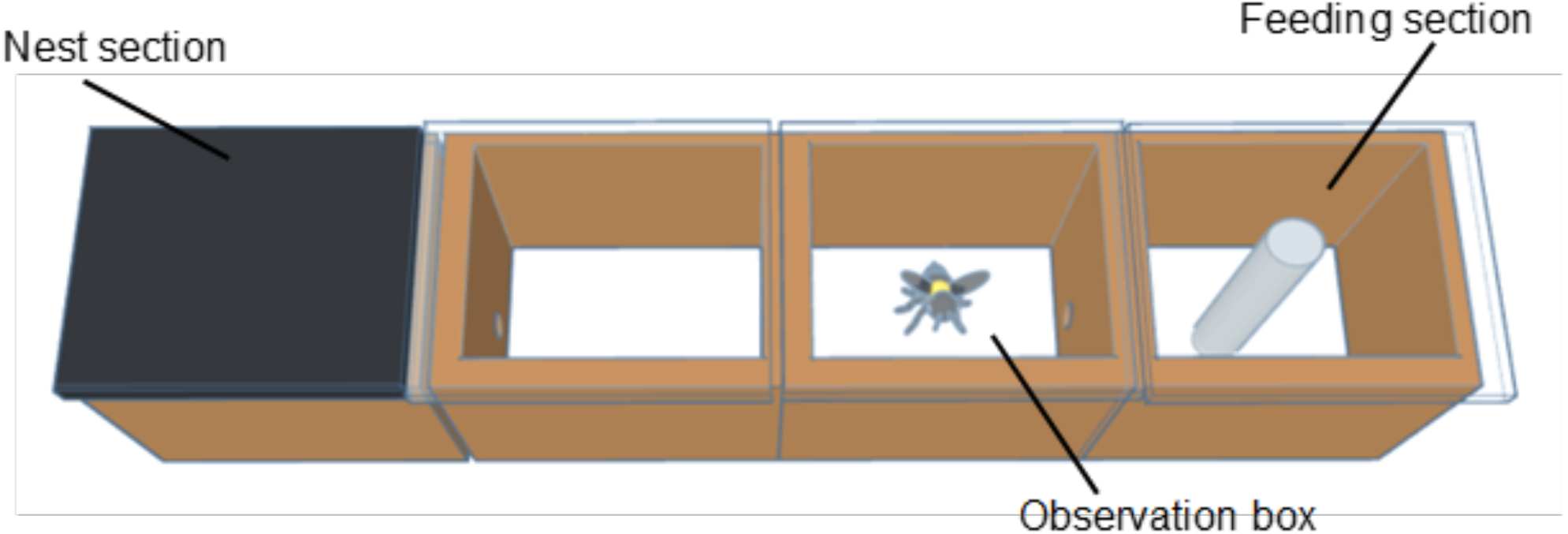
Housing and testing apparatus. A ventilated wooden box (56×16×11 cm) with four sections. The nest section was covered with plywood. The feeding section contained a feeder with *ab libitum* food. The observation box was adjacent to the feeding section.

### Treatments

For testing, we removed bees individually from the nest box by letting them walk onto metal forceps and placed them into a marking cage (Thorne, UK). A sponge in the marking cage was used to temporarily immobilize the bees to ensure precisely targeted noxious stimulation. A soldering iron (HAKKO FX-888D; Japan) was either heated to 65°C (noxious condition) or unheated (tactile condition), then touched onto the right or left antenna (counterbalanced across bees) for five seconds. We chose this method of noxious stimulation based on how stimulation of a honeybee’s (*Apis mellifera)* antenna with a 65°C heat probe causes consistent sting extension reflexes [44] (a defense reflex seen in response to noxious stimuli [45]). Thirty bees were touched with the noxiously-heated probe (noxiously-stimulated; N = 30); 28 were touched with the control unheated probe (tactilely-stimulated; N = 28); and 24 were put in the marking cage but not touched with a probe (control: N = 24). No bees were excluded from the analysis. We used an RST Soldering Iron Tip Thermometer 191 (YWBL-WH; China) to test the temperature of the soldering iron. After the treatment, bees were immediately placed in the observation box and filmed with an iPhone 8 (Apple; USA) for 25 minutes. We sealed the holes between boxes during the experiment, so bees were confined to the observation box (14×16×11 cm). We sexed each bee visually from the videos, based on the presence (in females) or absence (in males) of a black abdomen tip. There were 40 females and 18 males; sex was then accounted for in the statistical analysis.

### Behavioural analysis

Four treatment-blind coders recorded the grooming behaviour displayed in the 25-minute videos using BORIS behavioural analysis software (BORIS, version 7.9.15; Italy). Grooming was defined as ‘the right or left front, middle, or hind leg moves over the left or right antenna either in one direction or in a repeated back and forth motion’. To measure inter-rater reliability, all four raters recorded grooming behaviour for two bees (corresponding to two 25-minute videos: one noxiously-stimulated bee and one tactilely-stimulated bee). Because the rating scale was continuous, we calculated the intra-class correlation coefficient. The correlation compared the total grooming duration of the right and left antenna across the four raters. The coefficient was 0.86, on a scale of 0-1, indicating a ‘good’ reliability [46].

### Statistical analysis

We analysed the data in R (R Core Team, Cran-r-project, Vienna, Austria, version 2022.12.0+353), using generalised linear mixed effect models (GLMMs; packages: ‘lme4’ (Bates et al., 2015) and ‘car’ (Fox et al., 2021)) and Wilcoxon tests. We checked model assumptions using histograms and ‘Q-Q plots’, and corrected for multiple testing using the Holm-Bonferroni correction [48]. We considered p < 0.05 significant.

To test for a difference between the grooming duration on the touched versus untouched antenna in noxiously-stimulated and tactilely-stimulated bees, we ran a GLMM. The response variable was the duration of antennal grooming for each antenna per bee. The fixed effects were stimulation type (noxious or tactile), whether the antenna was touched or untouched, the sex of the bee and their interaction. The random effect was the bee identity. We ran this model for the whole observation period (25 minutes), as well as individual time bins 0-1, 0-2, 0-3, 0-4 0-5, 6-10, 11-15, 16-20 and 21-25 minutes. We tested the individual time bins because some previous invertebrate studies have only detected self-grooming within the first few minutes after stimulation [21–23]. The only time bin with a significant interaction effect (after applying the Holm-Bonferroni correction for multiple testing) was 0-2 minutes, so this is the only time bin we ran the other GLMM and Wilcoxon tests on (described below).

We used unpaired two-sample Wilcoxon tests (as our data did not meet the criteria for parametric analysis) to test the difference between the grooming durations on the touched or untouched antenna in the tactile and noxious treatment groups in the first two minutes after stimulation.

To test for a difference between grooming durations on either the touched or untouched antenna in the noxiously-stimulated and tactilely-stimulated bees, and the mean grooming duration for both antennae in bees in the control condition in the first two minutes, we ran another GLMM. The response variable was either the duration of touched antennal grooming per bee or the duration of untouched antennal grooming per bee (for control bees, the mean grooming on one antenna was used, because neither antenna was touched in this condition). The fixed effects were the stimulation type (noxious, tactile, control) and the sex of the bee. The random effect was bee identity.

## Results

We first tested whether there was a difference between grooming durations on the touched and untouched antennae, and, if so, whether this difference was larger when the probe was noxiously heated. For the whole 25-minute observation period, bees groomed their touched antenna significantly more (touched: 18.11±26.79 seconds; untouched: 2.22±3.57 seconds; t_5792_ = 5.922; p < 0.001; N = 40), regardless of whether the stimulation was noxious or non-noxious tactile (no significant effect: t_5792_ = 0.056, p = 0.955; N = 40; no significant interaction: t_5792_ = -0.224, p = 0.822; N = 40; Figure 2). Therefore, over the 25 minutes, grooming was directed towards the touched antenna, but not the noxiously-stimulated antenna specifically.

**Figure 2.**
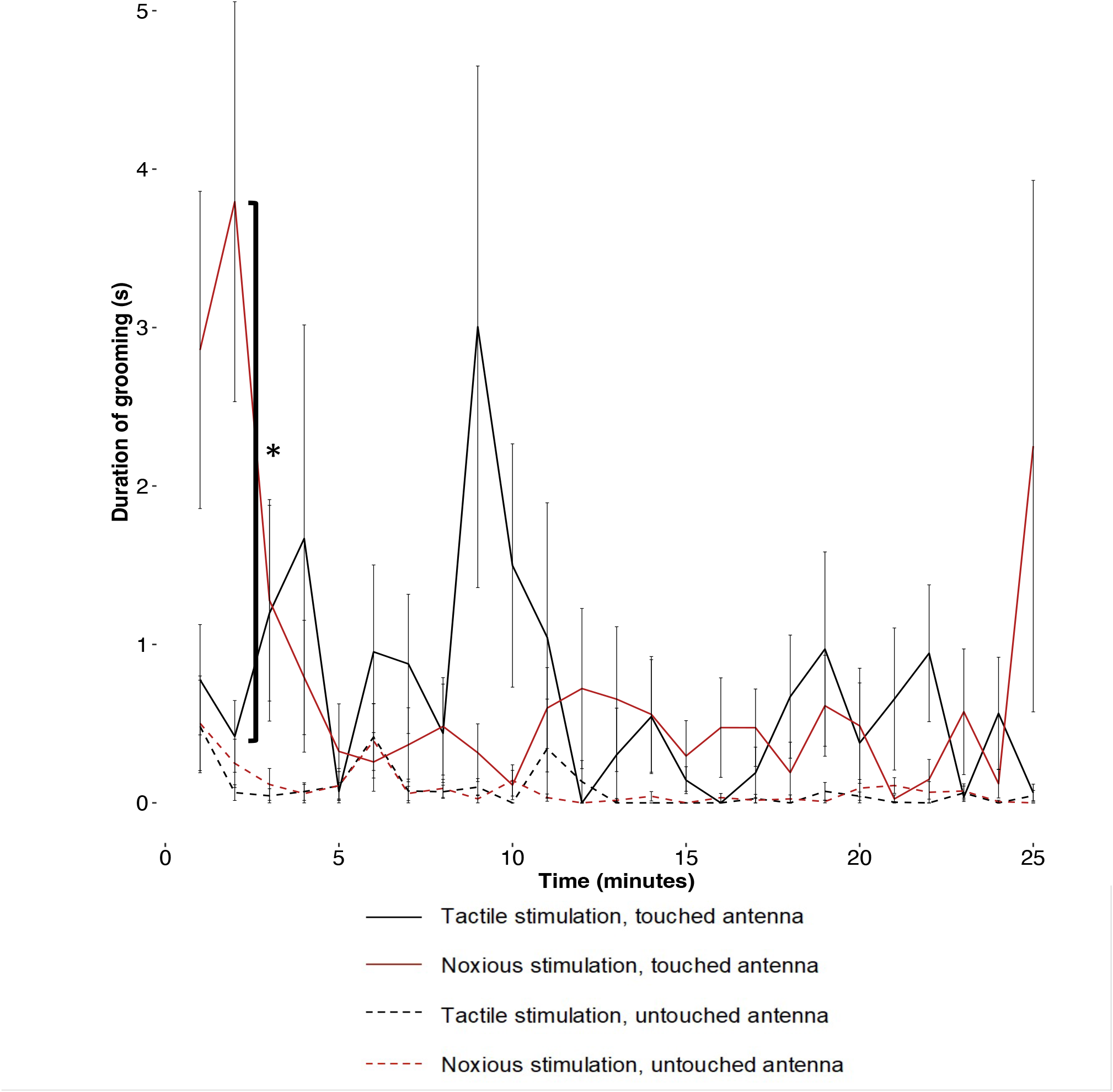
Duration of grooming for the untouched and touched antenna per each minute after noxious or tactile stimulation. P-values < 0.001 are represented by ‘*’; non-significant p-values are not represented in this graph.

We also observed a significant interaction effect of sex on the total grooming duration over the 25 minutes, with females grooming their touched antenna (and not their untouched antenna) significantly longer than males (females: N = 40; touched antenna: 22.89±30.29 seconds; untouched antenna: 2.60± 3.78 seconds; males: N = 18: touched antenna: 7.59±11.33 seconds; untouched antenna: 1.36±2.96 seconds; t_5792_ = -2.665; p < 0.01).

However, in the 0-2 minute time bin (the only time bin with a significant p-value after applying the Holm-Bonferroni correction), bees groomed the touched antenna more than the untouched antenna when the touch was noxious (significant interaction: t_459_ = 3.069, p < 0.005; N = 40). This result was clarified by Wilcoxon tests: in this time bin, noxiously-stimulated bees groomed their touched antenna (6.65±8.8 seconds) significantly more than their untouched antenna (0.75±1.95 seconds; W = 249.5, p < 0.001; N = 30; Figure 3). By contrast, for tactilely-stimulated bees, there was no difference in grooming between the touched antenna (1.19±2.23) and the untouched antenna (0.55±1.57 seconds; W = 324, p = 0.159; N = 28; Figure 3). There was no difference in the antennal grooming durations for male and female bees (t_459_ = -0.851, p = 0.395; N = 40).

**Figure 3.**
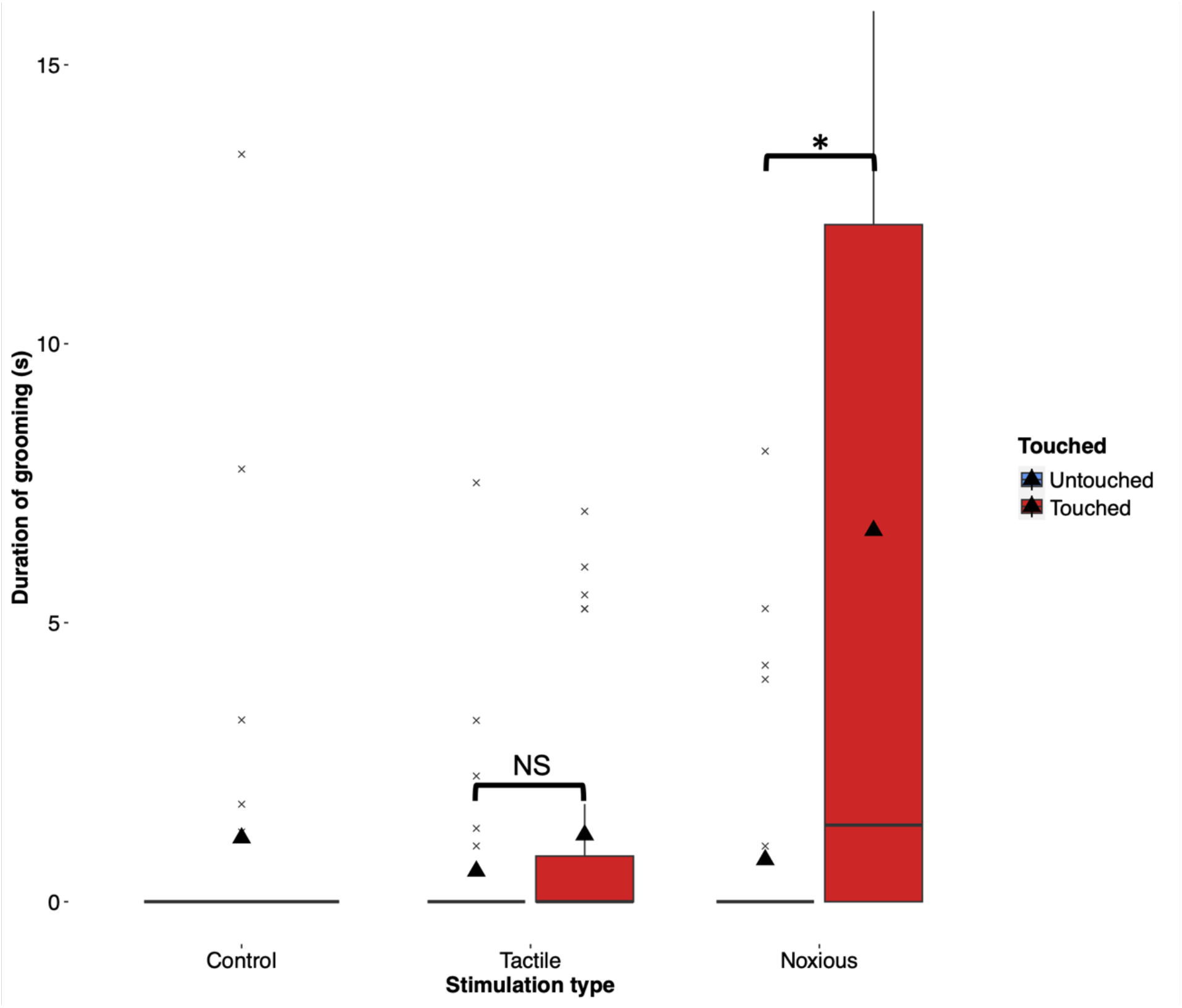
Duration of grooming each antenna for each treatment group. Box plot boundaries indicate the 25th and 75th percentiles; the whiskers indicate the minimum and maximum values within 1.5 times the interquartile range. Crosses indicate values outside this range (boxplot outliers); triangles indicate the mean; lines indicate the median. P-values < 0.001 are represented by ‘*’, and p-values > 0.1 are represented by ‘NS’.

We then tested whether the duration of antennal grooming was greater for either the noxiously-stimulated or the tactilely-stimulated bees compared to the control bees. Noxiously-stimulated bees groomed their touched, and not their untouched, antenna for longer than the control bees groomed either antenna (touched: 2.85±5.48 seconds; t_75_ = 2.55, p = 0.0127; N = 54; untouched: 0.50±1.64; t_75_ = -0.318, p = 0.752; N = 54; either antenna: 0.57±2.14; Figure 3). There was no significant effect of sex on either the grooming in touched or untouched conditions (touched: t_75_ = - 0.111, p = 9.117; N = 54; untouched: t_75_ = -1.493; p = 0.140; N = 54). By contrast, tactilely-stimulated bees did not groom either their touched or untouched antennae significantly more than the control bees groomed either antenna (touched: 0.77±1.84; t_73_ = -0.404, p = 0.689; N = 52; untouched: 0.48±1.54; t_73_ = 0.228, p = 0.821; N = 52; Figure 3; either antenna: 0.57±2.14). There was no significant effect of sex on either the grooming in touched or untouched conditions (touched: t_75_ = - 0.127, p = 0.210; N = 52; untouched: t_75_ = -1.875; p = 0.065; N = 52). Similarly, noxiously-stimulated bees groomed significantly more than the tactilely-stimulated bees on the touched antenna (t_83_ = 2.885, p < 0.005; N = 40; Figure 3), but not on the untouched antenna (t_83_ = 0.647, p = 0.519; N = 40; Figure 3). There was also no significant effect of sex on either the grooming in touched or untouched conditions (touched: t_83_ = 0.253, p = 0.800; N = 40; untouched: t_83_ = -1.273; p = 0.207; N = 40).

## Discussion

No previous studies have experimentally tested and quantifiably measured whether insects groom noxiously stimulated sites. This lack of evidence, as well as anecdotal reports of insects not demonstrating self-protective behaviour in other ways, has been used to argue that insects do not experience pain [30–32]. Our results provide clear, quantitative evidence of self-protective behaviour in insects.

In the first two minutes after stimulation, bees targeted grooming towards a noxiously-stimulated antenna, but not towards a tactilely-stimulated one. Noxiously-stimulated bees groomed their touched antenna more than their untouched antenna, more than tactilely-stimulated bees groomed their touched antenna, and more than control bees groomed either antenna. The same results were not found in tactilely-stimulated bees. These results were also only found in the first two minutes. Although there is an increase in grooming duration after noxious stimulation compared to tactile stimulation in the first minute (Figure 2), this increase is not significant after correction for multiple comparisons. This could be explained by the use of the Holm-Bonferroni correction, which has a high risk of false negatives [49]. There was a significant effect of sex on grooming in the first 25 minutes, with females grooming their touched antenna more than males. This is a potentially interesting result that could pave the way for some future research into sex differences in insects’ nociceptive responses. In this study, however, it does not affect the main result, so will not be discussed further.

Our results may hint at a potential control mechanism for nociception in insects. In mammals, rubbing a noxiously-stimulated site activates A-beta fibers [50]. The gate control theory posits that these fibers activate inhibitory interneurons in the dorsal horn of the spinal cord, which can inhibit the nociceptive signal’s progression to the brain [51,52]. This gated control means that ‘rubbing it better’ reduces nociceptive processing [12,13]. Self-grooming bumblebees may activate a similar mechanism. In honeybee antennae, thermo-sensory neurons detect the nociceptive stimulus and carry the information to the antennal lobe [53], and then, possibly, to a nociceptive thermal center in the brain [44]. The ‘gate’ could involve activating campaniform sensilla neurons in the antenna, although it is, so far, unclear how this might inhibit nociceptive processing. Further, some human experiments suggest that self-touch reduces the conscious aspect of pain, rather than just nociceptive processing [14,15]. Therefore, in particular, because of our results and other accumulating evidence for pain in insects, another mechanistic possibility is that grooming acts as a control mechanism for pain in bumblebees. Again, though, it is currently unclear how this might happen in insects.

Directed grooming of the noxiously-stimulated antenna only occurred in the 0-2 minute time bin. This timing is consistent with some studies on other invertebrates, which describe self-grooming in the first few minutes after noxious stimulation (17– 19). A reason for this timing might be that the nociceptive processing of the heat stimulation ceased after around two minutes; this would likely depend on the intensity of the noxious stimulus used. An association between grooming and the cessation or onset of nociceptive processing has been previously seen in mice, in response to formalin injection. There is an acute grooming phase, which apparently relates to the injection itself and lasts 3 minutes, then no grooming is seen for another 3 minutes, followed by a tonic phase that is longer-lasting and appears to correspond to formalin’s inflammatory effects [54–56]. By analogy, we suggest that, in our study, the first two minutes corresponded to an acute phase of grooming in response to the noxious heat stimulation. Based on this evidence, future research should investigate the neural processing of noxious heat stimulation in insects, and how the temporal characteristics of the grooming compare to the neural processing.

The experiment contained multiple novel and/or potentially stressful experiences and environments for the bees. For example, the stimulation itself involved them climbing onto metal forceps, being lifted out of the nest box, and immobilized during the stimulation - all potential stressors. Further, bees were isolated from the nest and other colony members during testing, and their normal route back to the nest was blocked. This means that the grooming we observed with this set-up may only be a fraction of the bees’ natural response, when not under stress or in a novel environment. Stress and novel contexts reduce the expression of behaviours after noxious stimulation in insects (honeybees: 57), similarly to other taxa (humans: 58; rodents: 59, fish: 60, birds: 61 and snails: 62). In future experiments, observing bees in the nest post-stimulation may reflect a more naturalistic behaviour.

Our study alone does not prove pain in insects, as there are multiple criteria for the experience of pain [8,9]. However, evidence for many of these other criteria has been shown in insects (see 37 for review). Importantly, our results provide clear evidence for a strong indicator of pain, self-protective behaviour [8], and undermine the claim that insects do not feel pain because they fail to show this behaviour. It is important to note that similar self-protective behaviour has been considered to be an indicator of pain in many other taxa, including crustaceans (21; molluscs (24; 25; 26), rodents [18], birds (19) and fish [20]. Excluding bumblebees from the same interpretation would be logically inconsistent.

*This research is part of a project that has received funding from Queen Mary University of London; an Erasmus+ traineeship; and the European Research Council (ERC) under the European Union’s Horizon 2020 research and innovation programme, Grant/Award Number 851145*.□□

## Acknowledgements

We thank Jonathan Birch for his insightful comments on and discussion about this manuscript.

